# Characterization of cutaneous reflexes elicited from saphenous nerve stimulation

**DOI:** 10.1101/2025.10.30.685609

**Authors:** Russell Hardesty, Sebastian Rueda Parra, Jonathan Wolpaw, Jodi Brangaccio

## Abstract

Phantom limb pain (PLP) is a chronic neuropathic condition that affects amputees. Cutaneous reflexes (CRs), mediated by spinal interneuronal circuits, represent a potential pathway linking non-nociceptive input to nociceptive modulation that could be targeted with neuromodulatory techniques such as operant conditioning. The objective of this study was to develop and validate a novel method for eliciting CRs from proximal muscles via saphenous nerve stimulation to enable future applications in individuals with PLP. We recruited 14 healthy adults and elicited CRs in the rectus femoris muscle via transcutaneous stimulation of the saphenous nerve at multiple stimulation intensities below their pain thresholds. We evaluated the consistency and reliability of CR response latencies and magnitudes, comparing the symmetry between the bilaterally (e.g. right and left legs). We found that CRs could be reliably elicited 50-100ms post-stimulus using saphenous stimulation and that responses were reliable at stimulation intensities below pain thresholds. Furthermore, we found the CR responses were relatively symmetrical in our healthy adult population. This study is the first of its kind to show that proximal (vs distal) CRs could be consistently elicited through the saphenous nerve stimulation. These findings imply that proximal CRs may be a feasible target for neuromodulatory interventions such as operant conditioning paradigms in the future.

## Introduction

Phantom limb pain (PLP), a form of chronic neuropathic pain, is a phenomenon where people who have undergone a limb amputation develop painful (e.g., sharp, aching or burning) sensations perceived to be occurring in their former limb. Phantom limb pain can occur not only after amputation, but also in children with congenital limb deficiency (Melzack, 1997). The mechanisms of phantom limb pain are still debated, although some evidence suggests that it may arise due to cortical reorganization, over-sensitization of peripheral nerves, or misinterpretation of the lack of sensory feedback from the absent limb as pain (Melzack, 1990; D’Mello and Dickenson, 2008; Petersen et al., 2019; Felicetti et al., 2021). In addition to PLP, the residual limb frequently has increased sensitivity to non-nociceptive inputs, e.g., tactile stimuli (Ropero Peláez and Taniguchi, 2016; Eckert et al., 2022).

Cutaneous reflexes (CRs) elicited from non-nociceptive stimulation are generated from a complex network of spinal interneurons and play an important role in modulating muscle activity for a wide range of motor tasks, including locomotion, posture maintenance, and the pain withdrawal reflex (Zehr et al., 2001; Viseux et al., 2019; Felicetti et al., 2021). Cutaneous reflexes are not fixed responses; rather, they are considered functionally adaptive and dynamically modulate their expression based on the demands and context of the motor task. They are modulated by supraspinal inputs and can be either excitatory or inhibitory depending on factors such as gait phase, limb position, or voluntary intent. For example, during walking, cutaneous reflexes may be facilitatory or inhibitory based on the gait phase, the type of walking surface, and contextual-based anticipatory adjustments (Duysens et al., 1996; Zehr and Stein, 1999; Zehr and Haridas, 2003). Additionally, cutaneous reflexes may have a critical role in spinal processing of nociceptive stimuli. In 1965, Melzack and Wall first proposed the gate control theory of pain which posited that Aδ and C fibers responsible for transmitting nociceptive stimuli could be modulated by the larger Aꞵ fibers which responded to non-nociceptive stimuli (Melzack and Wall, 1965). According to this model of pain, both nociceptive and non-nociceptive stimuli are integrated in the substantia gelatinosa – under central control of cortical processes, effectively “closing the gate” – in an antagonistic role with one another, i.e. increased non-nociceptive input will inhibit transmission of nociceptive stimuli. Since its initial formulation, the gate control theory has evolved to acknowledge a more distributed pain network throughout the nervous system, culminating in the “neuromatrix” theory (Melzack, 1999). However, Aꞵ fibers – the fibers responsible for the cutaneous reflex – are still recognized as a modulator and transmitter of nociceptive stimuli (D’Mello and Dickenson, 2008). Disruption of these fibers and reflexes due to injury or illness can lead to faulty or inappropriate nociceptive responses. Thus, approaches that target the cutaneous reflex, such as operant conditioning protocols, are plausible interventions for chronic pain (Phipps and Thompson, 2024, 2025).

The objective of this study was to characterize proximal cutaneous reflexes evoked by saphenous stimulation to assess whether they may be a feasible target for operant conditioning protocols aimed at treating PLP. Previous studies have primarily elicited cutaneous reflexes almost exclusively using distal cutaneous nerve stimulation, e.g., sural nerve (Brooke et al., 1999; Zehr et al., 2001; Phipps and Thompson, 2024). This would be infeasible for most lower limb amputees. Moreover, the methods used across studies vary considerably, without standardized protocols and differences in stimulation parameters. Therefore, we developed a protocol to elicit cutaneous reflexes in more proximal muscles, such as *rectus femoris*, by stimulating the saphenous nerve on the medial portion of the knee. We then characterized the resulting CRs in 14 healthy controls. We evaluated CR latency and magnitude at five stimulation intensities and calculated coefficients of variation and intraclass correlation coefficients to assess CR reliability. Furthermore, we examined interlimb asymmetries in cutaneous reflexes among healthy participants, providing a normalized reference for assessing potential deviations in individuals with traumatic amputation.

## Methods

### Ethics statement

All procedures were approved by the Institutional Review Board (IRB) of the Stratton VA Medical Center consistent with the standards of the *Declaration of Helsinki* (Protocol #: 1584762). Participants provided written informed consent to participate in the study.

### Human participants

Fourteen healthy individuals (5 females, 9 males; ages 49.3 ± 14.6 years (mean ± standard deviation), ranging 29 to 72 years old) participated in the study after providing written informed consent. The study was performed at the Albany Stratton VA Medical Center, Albany, NY and approved by the Albany Samuel Stratton Veterans Affairs Medical Center (SSVAMC) IRB. Inclusion criteria included: (1) ≥18 years old; (2) no known history of neurological disease; (3) no significant cardiac history; and (4) measurable cutaneous reflex by saphenous nerve stimulation.

### Experimental Design

Adapting and modifying from the methodology utilized with the H-Reflex Operant Conditioning (HROC) protocol also used in our lab, we elicited cutaneous reflexes (CRs) proximally in leg muscles stimulating the saphenous nerve (Thompson et al., 2013, 2019, 2022; Makihara et al., 2014; Kim et al., 2023).

### EMG

At the beginning of each session, both legs were cleaned with alcohol and gently abraded prior to the application of EMG recording and stimulating electrodes. For recording EMG, surface Ag-AgCl electrodes were placed longitudinally in bipolar fashion over the stimulated leg’s Rectus Femoris (RF), Vastus Lateralis (VL), Vastus Medialis (VM), Tibialis Anterior (TA), Medial Gastrocnemius (MG), the ground was placed on the patella and the non-stimulated leg’s RF MG, TA muscles approximately 3 cm apart center-to-center. (Makihara et al., 2014; Thompson et al., 2013, 2009). EMG was amplified (gain=500) and bandpass filtered (10–1000 Hz) (AMT-8 amplifier, Bortec Biomedical Ltd., Calgary, Alberta, Canada), and then digitized (3200 Hz) and stored using our evoked potential operant conditioning software (EPOCS) (Hill et al., 2022).

### Electrical stimulation

Stimulation electrodes were applied to the medial knee region targeting the saphenous nerve through surface Ag-AgCl electrodes (2.2 × 2.2 cm) with the cathode along the saphenous nerve medial to the patella and around the level of the joint line, the anode 4 cm distally. Electrode position was then optimized to minimize the stimulation current required to elicit a threshold HR, while maximizing HR size and participant comfort. CRs were primarily analyzed in the rectus femoris (RF) muscle (**Figure 2a**), which was the actively monitored channel used to trigger stimulation. The RF CR was elicited by transcutaneous electrical stimulation (DS8R stimulator, Digitimer Ltd, Welwyn Garden City, UK) using a train of 15 biphasic pulses (0.5 ms pulse width, positive polarity), intratrain interval of 1 ms and intertrain interval of 2 s ±20%. Therefore, each train took ∼ 30 ms with 2 s ± 20% in between trains. Reflexes were recorded bilaterally, with testing always beginning on the right leg (self-reported dominant leg in 14/14 participants).

### CR Elicitation Protocol

During each trial, participants maintained a modest level of RF (channel 1) electromyographic (EMG) activity within a predefined window (typically 5–18 µV) through a sustained isotonic contraction for 200 ms. Additionally, to limit lower limb involvement, the Tibialis Anterior (TA) was also monitored and controlled to be no greater than 0-15 µV. Stimulation was only delivered when EMG criteria was met for both channels and maintained for 200 ms. Participants received real-time visual feedback via a screen positioned in front of them: a colored bar displayed muscle activity status, turning red when EMG activity fell outside the target windows and green when both conditions were satisfied (**Figure 1B**). Stimulation consisted of a train of 15 biphasic pulses (0.5-ms pulse width, positive polarity), intratrain interval of 1 ms and intertrain interval of 2 s ±20%.

**Figure 1:**
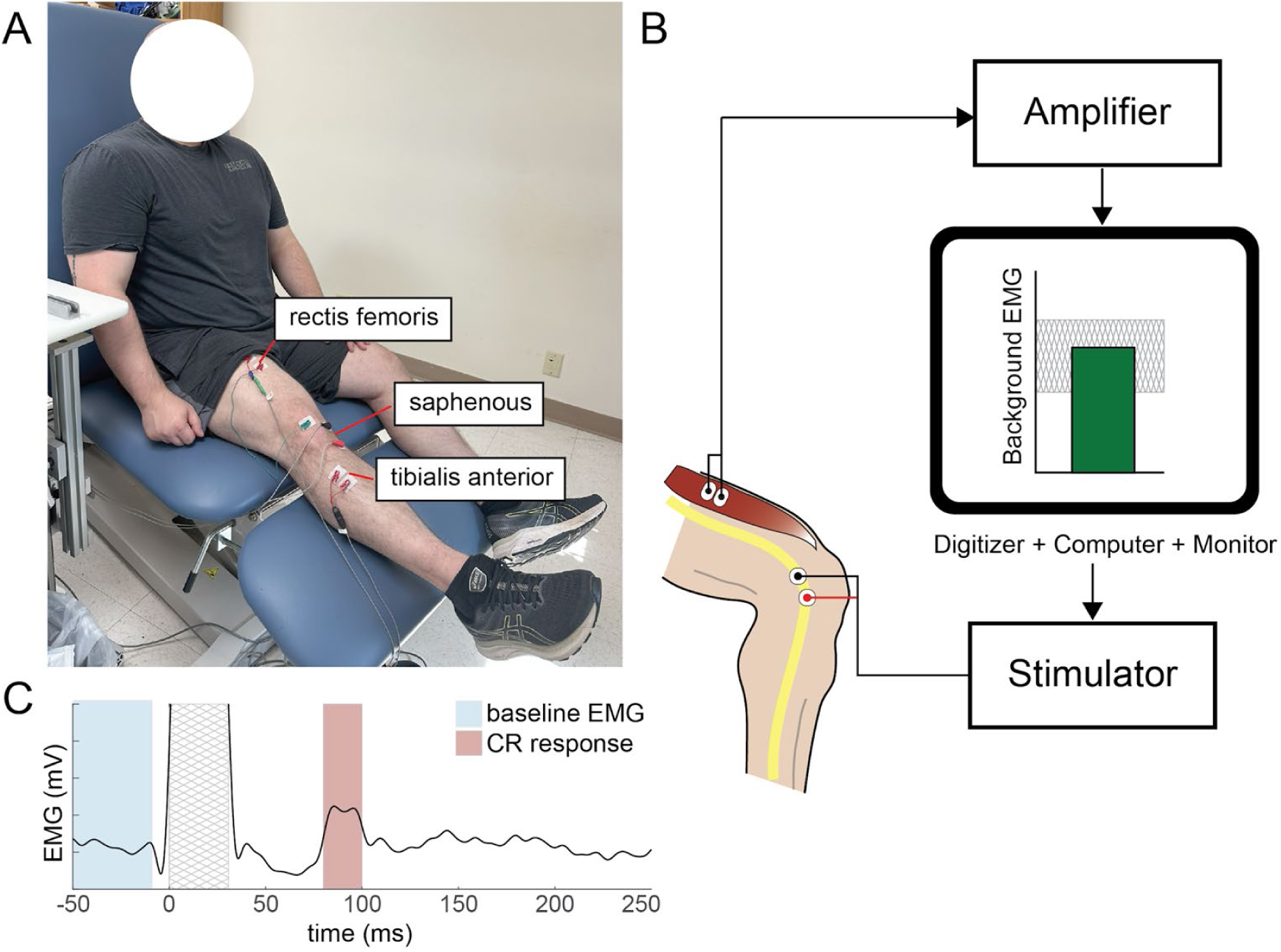
(A) Experimental setup with stimulating and recording electrode locations. (B) Schematic of EPOCS system used to trigger stimulation only when background EMG is within desired range. (C) Example of cutaneous reflex (CR) response (red region) and the baseline pre-stimulus EMG activity (blue region).

**Figure 2:**
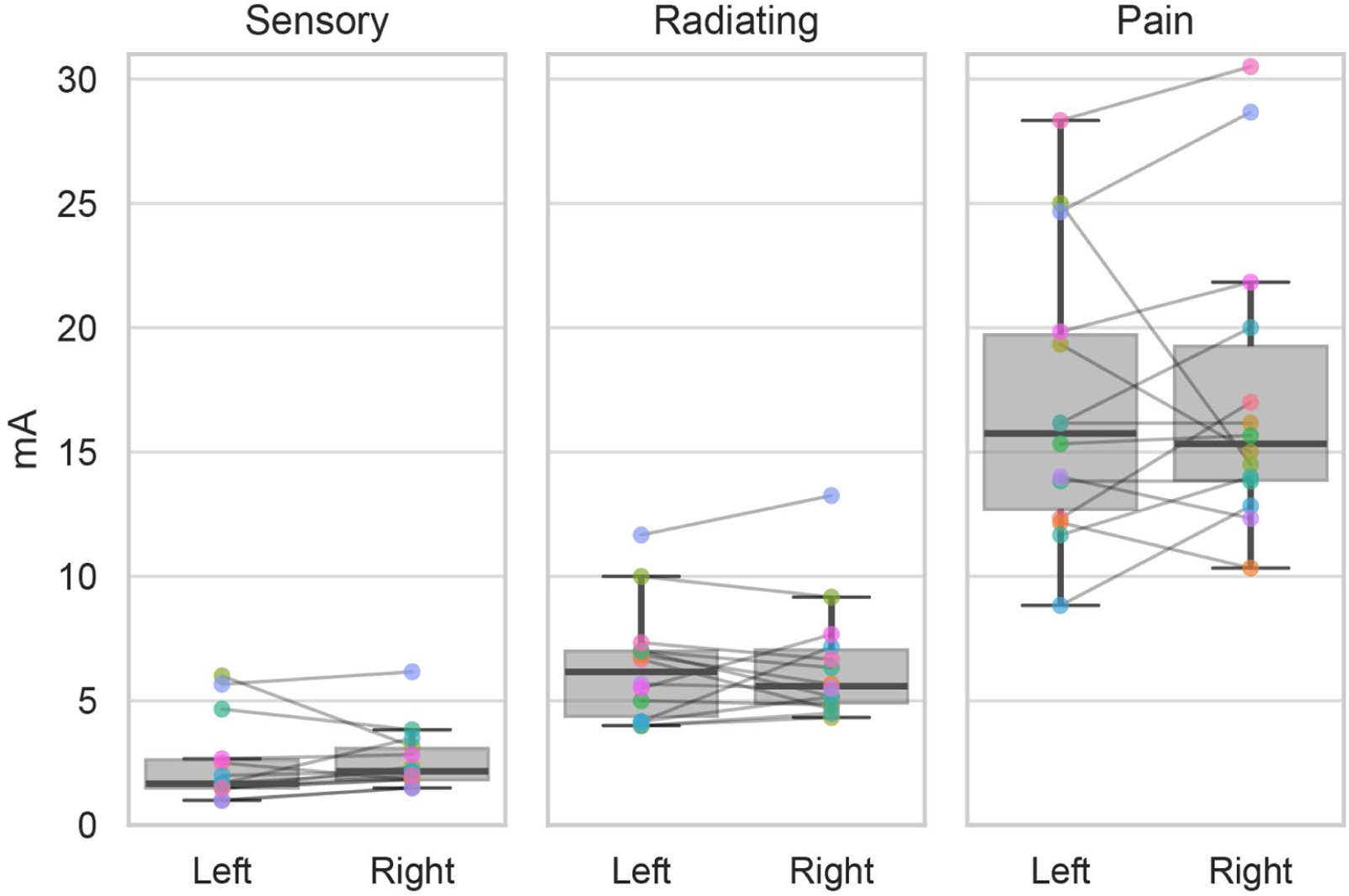
Sensory, radiating, and pain thresholds for each participant bilaterally. Data points represent a single participant with lines connecting the threshold for the left and right legs of the same individual.

Stimulation intensity was normalized for each leg based on three perceptual thresholds: (1) detection (first noticeable sensation of stimulation, also known as sensory threshold (ST)), (2) radiating threshold (RT) (sensation extending beyond the stimulation site), and (3) discomfort (pain threshold (PT)) (**Figure 2.b.**). Thresholds were measured three times, by gradually increasing the stimulation current in steps of 0.2-0.5 mA, participants were instructed to report: 1) when they first felt the electrical stimulation (ST); 2) when they perceived the stimulation sensation spread outside and away from the stimulation pads in any direction (RT), also noting the extent and direction of spread; and 3) when the stimulation became uncomfortable (PT). Each of these thresholds were averaged to determine stimulation levels using Eq 1:

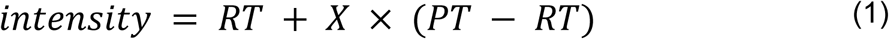

Here *X* is the unitless stimulation intensity normalized to RT and PT. For example, if RT = 5mA and PT=10mA, then a relative stimulation intensity of 50% (X=0.5) would result in a stimulation intensity of 7.5mA. Five relative stimulation levels were selected, defined as percentages of the difference between the radiating and pain thresholds: −25%, 25%, 50%, 75%, and 100%. For each level, 30 repetitions were collected, starting with the self-reported dominant (right) limb.

### Signal Processing

All signal processing and analyses were performed using custom Python scripts. EMG signals were low-pass filtered (fc = 100 Hz) using a Butterworth filter and full-wave rectified. Data was epoched relative to stimulus onset, from –50ms to +200ms. To quantify CR response, we calculated the mean magnitude of the rectified signal within the time window of interest. CR responses were normalized to baseline EMG activity which was calculated as the mean rectified activity in the period –50 to –10ms prior to the stimulation (**Figure 1C**).

### Symmetry

We computed a symmetry index by calculating the difference of the mean rectified CR response in the left and right limbs normalized to the total of the two responses as follows:

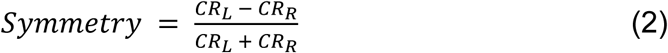

Here CRL and CRR are the mean rectified magnitude of the cutaneous reflex for the left and right leg, respectively.

### Statistical analysis

Sensory, radiating, and pain thresholds were compared between limbs using paired t-tests. Effect sizes were quantified using Cohen’s d. To assess within– and between-participant reliability, we calculated coefficients of variation and two-way mixed, average-measures intraclass correlation coefficients (ICC3k) (Shrout and Fleiss, 1979). Independent t-tests were used to compare CR magnitudes with baseline EMG activity and to test whether symmetry indices differed significantly from zero. For all hypothesis tests, statistical significance was evaluated using an *a priori* alpha level of ɑ = 0.05.

## Results

### Sensory, radiating, and pain thresholds are symmetrical

This study sought to characterize proximal CR via the saphenous nerve in HC. To do this we, first, elicited CR responses via the saphenous nerve in 14 HC bilaterally. We determined sensory (ST), radiating (RT), and pain (PT) thresholds for each participant bilaterally. We compared ST, RT, and PT of the left and right leg using a paired t-test. We found no differences between limbs in ST(t-statistic = –0.464, p-value = 0.6505, Cohen’s d = –0.124), RT (t-statistic = –0.198, p-value = 0.8462, Cohen’s d = – 0.053, or PT (t-statistic = –0.328, p-value = 0.7483, Cohen’s d = –0.088).

### Identifying CR response latency

Cutaneous reflexes arise from the activation of multiple classes of sensory afferents with distinct conduction velocities, including Aβ, Aδ, and C fibers. Integration of these afferent inputs within the spinal cord is largely polysynaptic and often involves projections to multiple motor neuron pools, resulting in a nonsynchronous, temporally dispersed response that can be observed in full-wave rectified EMG. Previous studies of cutaneous reflexes in humans have primarily focused on distal lower-limb stimulation, such as the sural nerve, and have demonstrated both facilitatory (positive) and inhibitory (negative) EMG responses. Importantly, these responses are often task-dependent; for example, stimulation that elicits facilitation during standing may evoke inhibition during walking. Consistent with these findings, we observed that saphenous nerve stimulation also produced both facilitatory and inhibitory responses in rectus femoris EMG (Figure 3).

**Figure 3:**
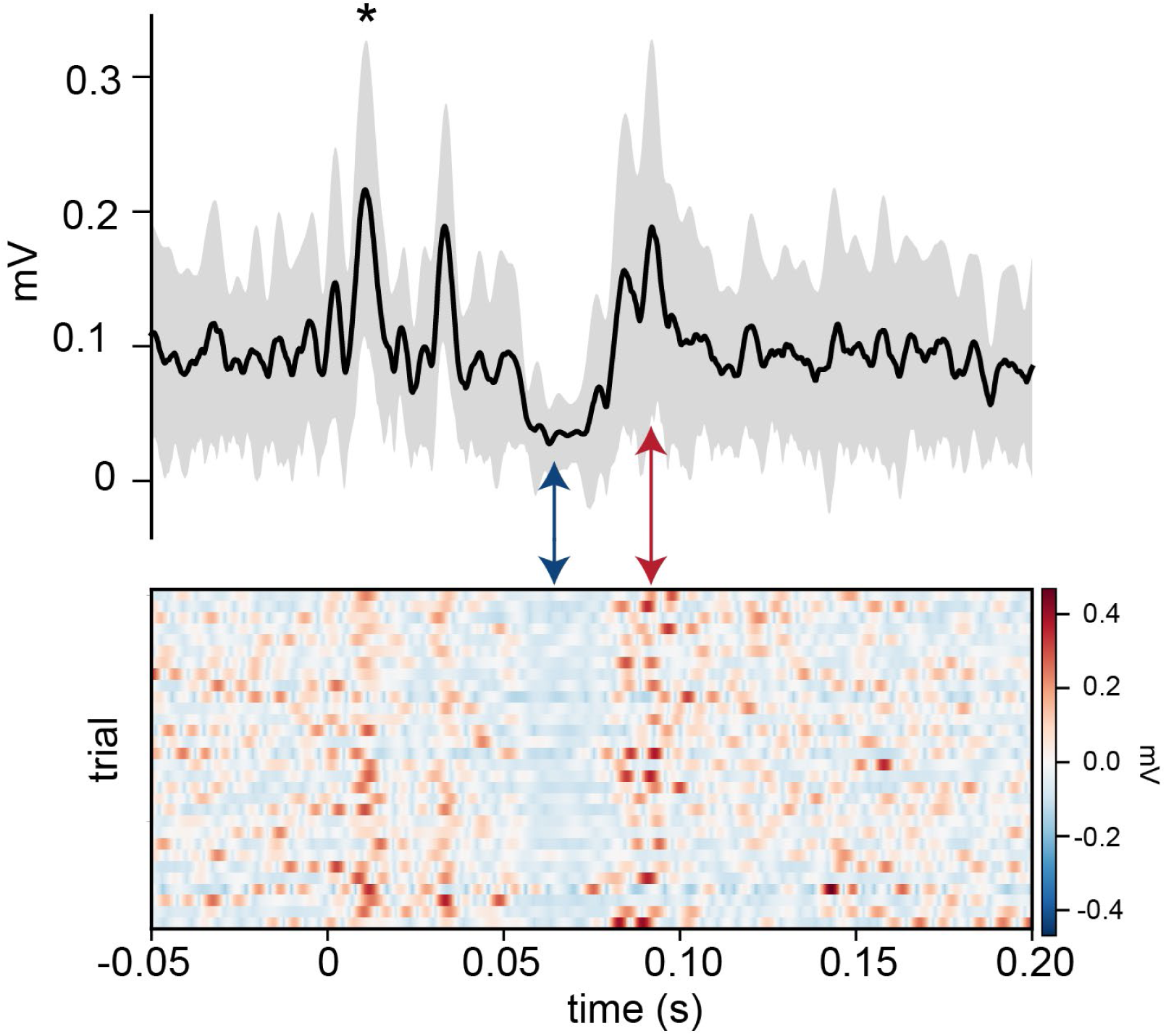
(Top) Example of the cutaneous reflex generated from stimulation of saphenous nerve for a single participant recorded from rectus femoris. The solid line denotes the mean across N=30 stimulations; the shaded region denotes the standard deviation across trials. Stimulation occurred at time 0 and stimulation artifact is marked with an asterisk (*). (Bottom) Each row of the heatmap shows an individual stimulation. Following stimulation EMG activity shows distinct periods of inhibition (blue arrow) and facilitation (red arrow).

Because CRs have not previously been studied during saphenous nerve stimulation, we made no *a priori* assumptions about response latencies or their consistency across participants. For reference, prior studies using sural nerve stimulation have categorized EMG responses as short (40–80 ms), moderate (80–120 ms), or long latency (120–150 ms). To evaluate the latency and consistency of CRs in our experiment, we constructed peristimulus time histograms (PSTHs) using participant-level mean responses at a stimulation intensity of 75% (Figure 4). We selected 75% because it was the highest intensity tested that remained below each participant’s pain threshold. The PSTH was computed by binning the EMG signal into 5-ms intervals and calculating the mean rectified EMG amplitude for each bin. We then compared each poststimulus bin to the prestimulus background EMG and defined a response as any bin where the EMG magnitude exceeded ±1.5 standard deviations of the background mean. This threshold was chosen as the lowest criterion that limited false positives to fewer than 5% in prestimulus bins. Finally, to assess cross-participant consistency, we summed the number of participants showing a response in each bin, allowing us to identify latencies where CRs were most reliably observed.

**Figure 4:**
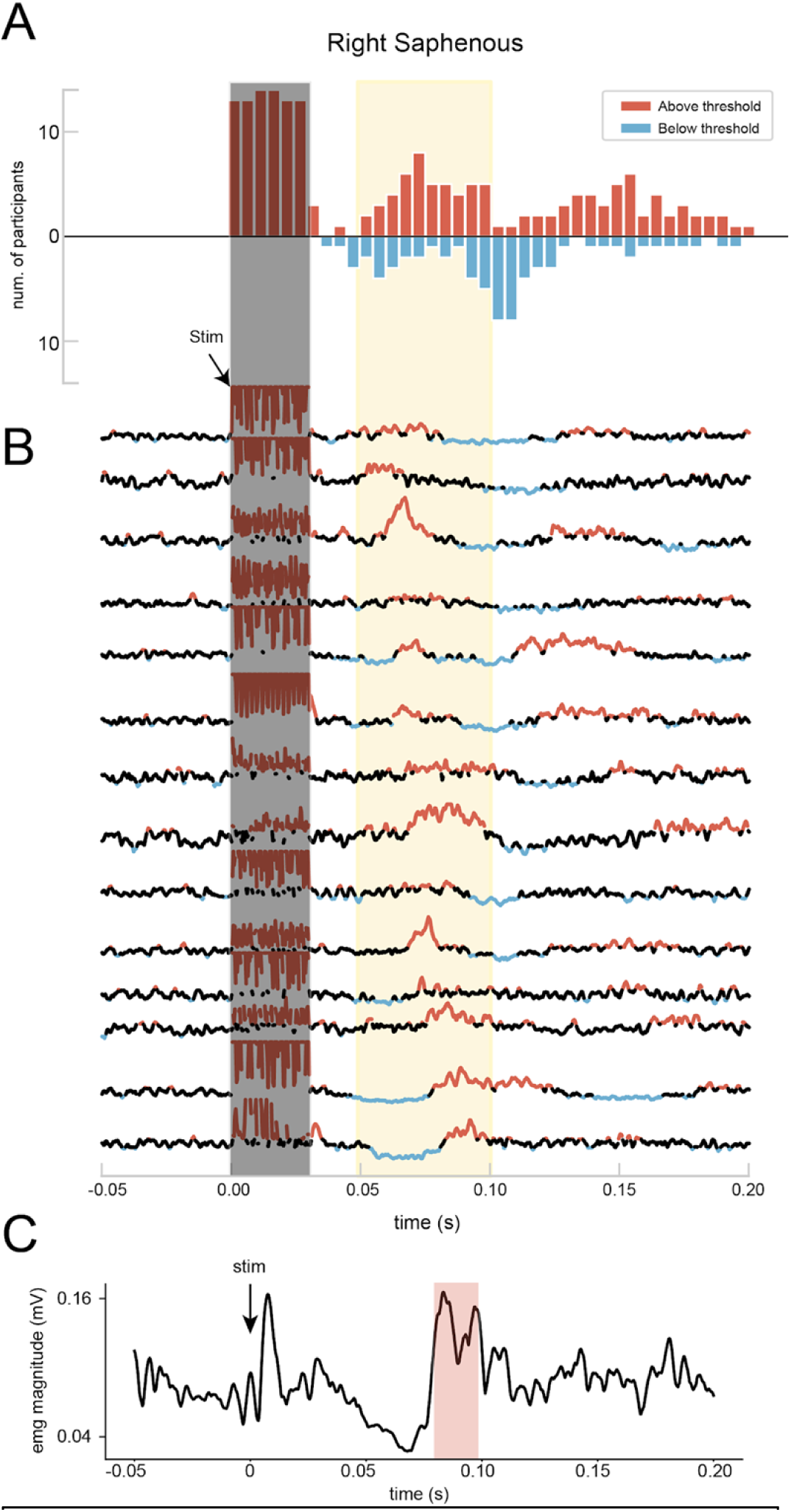
(A) PSTH of EMG activity greater (red) or less (blue) than 1.5 times the standard deviation. (B) Average response for each of N=14 participants with regions above and below the threshold denoted. (C) Example of automatic response detection.

We found that response latency and direction (facilitatory vs inhibitory) did differ across participants. However, the PSTH showed facilitatory responses in a majority of participants between 50-100ms (Figure 4A). Upon visual inspection of the participant mean responses in that window, we found that all participants demonstrated EMG activity greater than 1.5 standard deviations for a portion of the 50-100ms region, but that the specific latencies of each response did differ across participants within that window (Figure 4B). Therefore, we defined participant-specific time windows to quantify the CR response. We performed a continuous wavelet transform using a Morlet wavelet template to identify participant-specific boundaries for the CR response similar to methods used to automatically detect H-reflex waveforms (McKinnon et al., 2023) (Figure 4C). We quantified CR responses by calculating the mean rectified EMG magnitude within these participant-specific windows.

### Intra– and inter-participant CR consistency

To assess the intra– and inter-participant consistency of the quantified CR responses, we calculated the two-way mixed, average-measures intraclass correlation coefficient (i.e., ICC3k) and the coefficient of variation (CV) for each stimulation intensity and site (Table 2). Across all stimulation intensities, mean CV values ranged from 0.50 to 0.61, indicating moderate trial-to-trial variability within participants irrespective of stimulation intensity. In contrast, the ICC3k values increased with stimulation intensity, with the highest intensities (75% and 100%) showing good to excellent reliability (ICC3k = 0.77–0.86). These findings indicate that although individual trial responses are somewhat variable, the averaged CR responses are highly reliable at larger stimulation intensities.

**Table 1:**
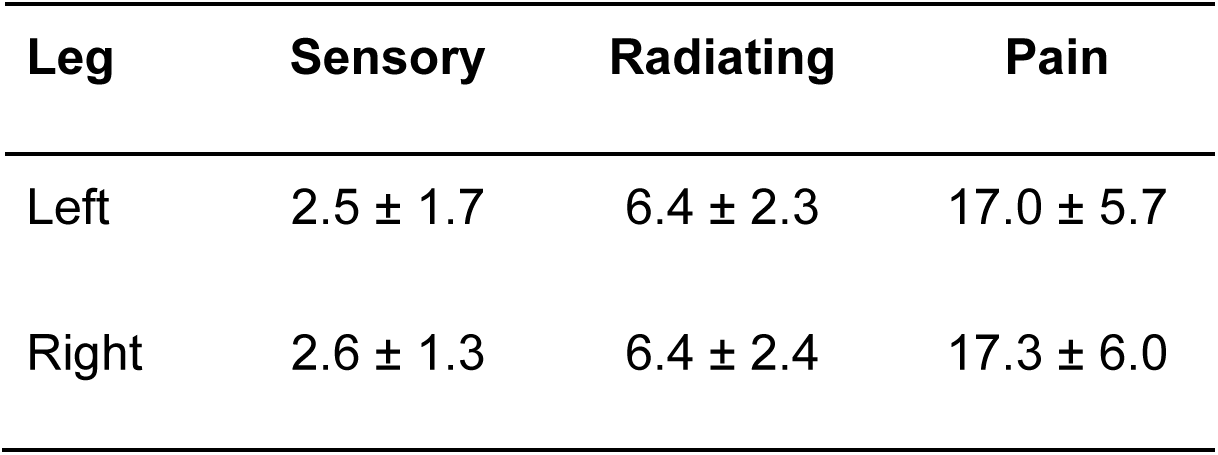
Mean threshold intensities (mA)

**Table 2:**
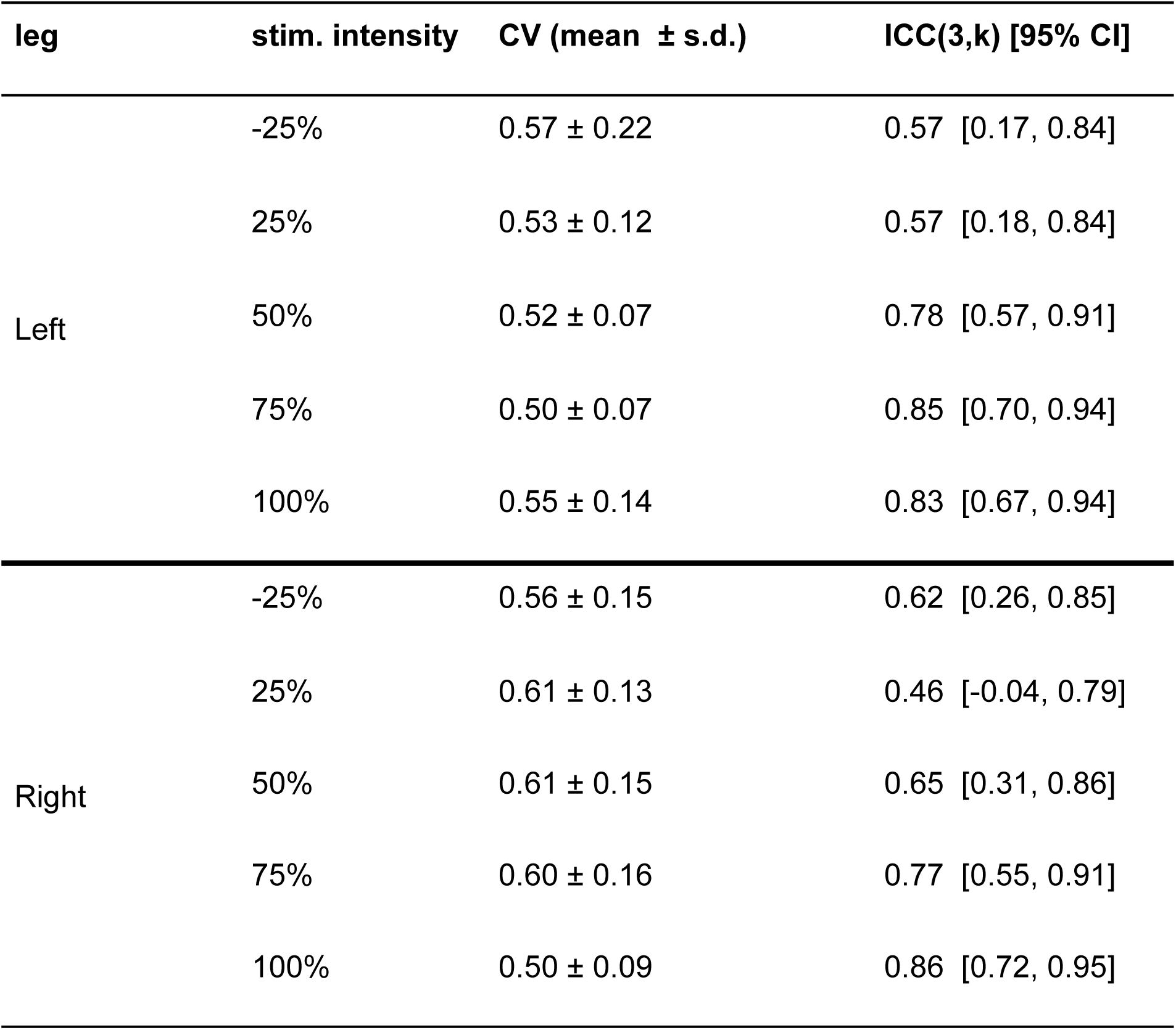
Inter– and intra-participant consistency.

| leg | stim. intensity | CV (mean ± s.d.) | ICC(3,k) [95% CI] |
| --- | --- | --- | --- |
| Left | -25% | 0.57 ± 0.22 | 0.57 [0.17, 0.84] |
| | 25% | 0.53 ± 0.12 | 0.57 [0.18, 0.84] |
| | 50% | 0.52 ± 0.07 | 0.78 [0.57, 0.91] |
| | 75% | 0.50 ± 0.07 | 0.85 [0.70, 0.94] |
| | 100% | 0.55 ± 0.14 | 0.83 [0.67, 0.94] |
| Right | -25% | 0.56 ± 0.15 | 0.62 [0.26, 0.85] |
| | 25% | 0.61 ± 0.13 | 0.46 [-0.04, 0.79] |
| | 50% | 0.61 ± 0.15 | 0.65 [0.31, 0.86] |
| | 75% | 0.60 ± 0.16 | 0.77 [0.55, 0.91] |
| | 100% | 0.50 ± 0.09 | 0.86 [0.72, 0.95] |

### CR magnitude increases with stimulation intensity

We examined the effect of stimulation intensity on cutaneous reflex (CR) magnitude by normalizing the mean rectified CR response to baseline EMG. Independent t-tests confirmed that normalized CR responses were significantly greater than baseline even at the lowest intensity tested (−25%) for both legs (Table 3). CR magnitude increased with stimulation intensity and reached an apparent plateau at approximately 75% intensity (Figure 5).

**Figure 5:**
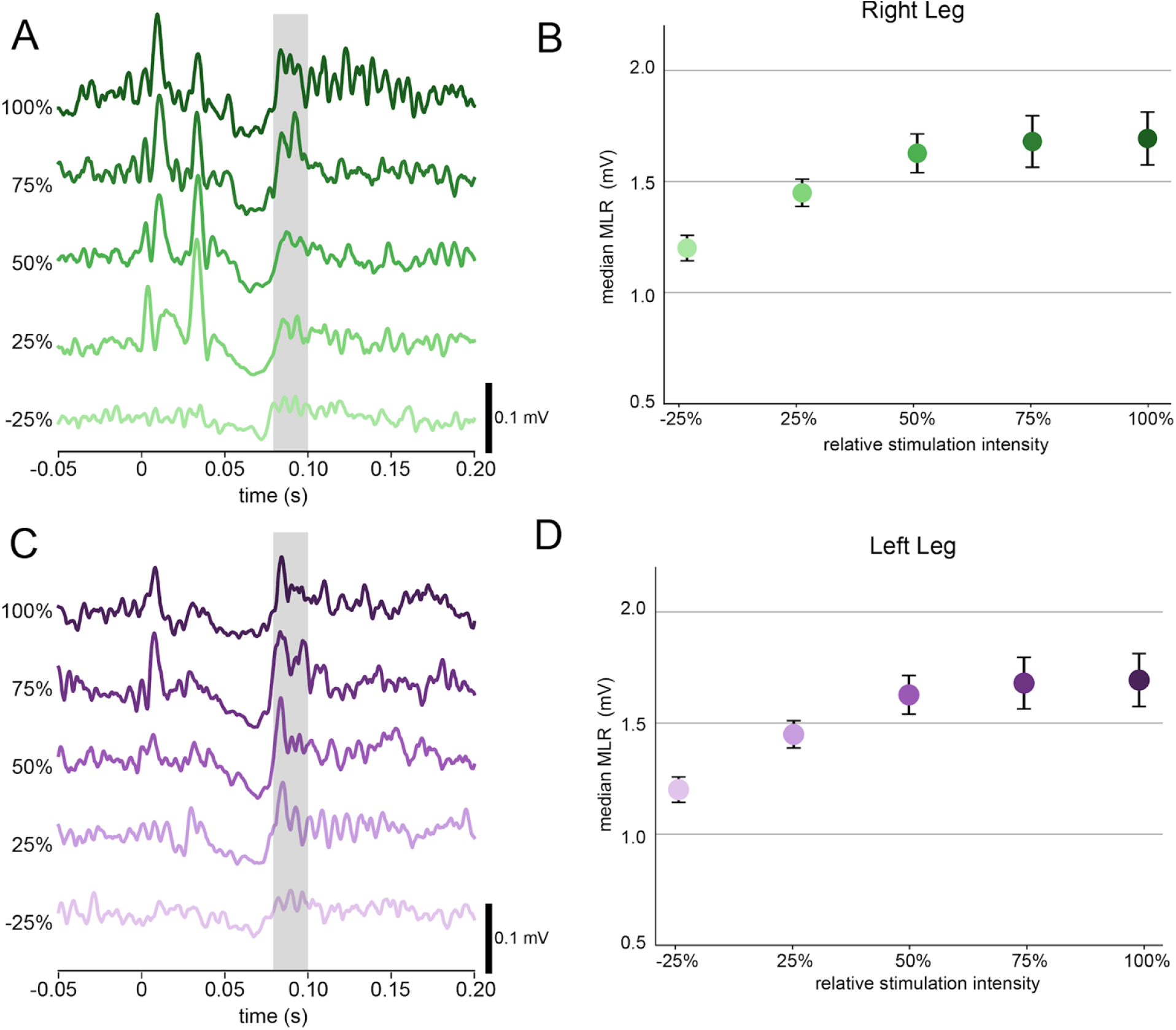
(Left) Mean cutaneous reflex responses for the right (A) and left (C) legs at increasing relative stimulation intensities. (Right) Quantification of MLR size across participants. Dots denote

Table 3:

**Table 3:**
| **leg** | **intensity** | **mean response [95% CI]** | **t-statistic** | **p-value** |
| --- | --- | --- | --- | --- |
| left | -25% | 1.20 [1.08, 1.32] | 3.50 | <0.01 |
|  | 25% | 1.45 [1.32, 1.58] | 7.31 | <0.001 |
|  | 50% | 1.63 [1.44, 1.82] | 7.20 | <0.001 |
|  | 75% | 1.72 [1.47, 1.97] | 5.86 | <0.001 |
|  | 100% | 1.69 [1.44, 1.95] | 5.82 | <0.001 |
| right | -25% | 1.24 [1.11, 1.37] | 4.06 | <0.01 |
|  | 25% | 1.31 [1.19, 1.43] | 5.60 | <0.001 |
|  | 50% | 1.48 [1.30, 1.66] | 5.83 | <0.001 |
|  | 75% | 1.58 [1.36, 1.80] | 5.67 | <0.001 |
|  | 100% | 1.59 [1.34, 1.83] | 5.10 | <0.001 |

### Left-Right Symmetry of CR

The long-term goal of characterizing CR responses elicited with saphenous stimulation was to evaluate whether CRs may be a feasible target for operant conditioning protocols in limb amputees. Therefore, we evaluated the CR responses between the left and right legs to determine presence and the extent of variability between legs in HCs. Across participants, CR magnitudes were largely symmetric, with most symmetry indices ≤ 0.1, except for two outliers. An independent t-test confirmed that the population symmetry index did not differ significantly from zero (t-statistic = –1.18, p = 0.26).

## Discussion

### Summary

The primary objective of this study was to characterize cutaneous reflexes evoked by saphenous nerve stimulation to assess CRs as a viable target for operant conditioning protocols. To address this goal, we (1) assessed the reliability of reflex latency and magnitude both within and across participants, (2) examined the relationship between reflex magnitude and stimulation intensity, and (3) evaluated left–right symmetry of reflex magnitudes. Across all participants, stimulation consistently elicited a facilitatory reflex response occurring 50–100 ms post-stimulation. Intraclass correlation coefficients (ICC3k) showed good reliability at stimulation intensities greater than 50%. Averaged reflex magnitudes increased systematically with increasing stimulation intensity for all participants bilaterally, reaching a plateau at approximately 75% of relative stimulation intensity. Finally, averaged reflex magnitudes were largely symmetrical between limbs, with interlimb differences <10% in most participants.

### CR latency, magnitude, and reliability

Cutaneous reflexes recorded via electromyography (EMG) have been studied primarily using sural nerve stimulation. These EMG responses are commonly classified according to their latency: short-latency responses (SLRs) occurring approximately 50– 80 ms post-stimulation, middle-latency responses (MLRs) occurring at 80–120 ms, and long-latency responses (LLRs) occurring after 120 ms. SLRs are thought to arise from fast-conducting, oligosynaptic Aβ afferent pathways within the spinal cord, whereas MLRs are believed to involve more polysynaptic mostly spinal but can occasionally receive supraspinal input. In contrast, LLRs are mediated primarily by supraspinal pathways, likely reflecting contributions from sensorimotor cortical processing (Bawa and McKenzie, 1981; Jenner and Stephens, 1982; Macefield et al., 1996; Schieppati et al., 1996). CR responses can be either facilitatory or inhibitory at these latencies and may change depending upon the task. For example, MLRs are often facilitatory during standing but may become inhibitory during walking (Phipps and Thompson, 2023). Because this study examined cutaneous reflex (CR) responses elicited by stimulation of the more proximal saphenous nerve, we did not make a priori assumptions regarding either the latency or direction of the responses. Instead, we observed both facilitatory and inhibitory EMG regions across participants within the 50–100 ms post-stimulation window (Figure 4).

This variability poses a challenge for quantifying CR magnitude, as the presence of both facilitatory and inhibitory components within a single predefined time window can result in signal cancellation and underestimation of response amplitude. To address this, we defined participant-specific quantification windows to more accurately capture individual response characteristics which would, in the context of operant conditioning, provide more reliable feedback to participants. Using this individualized approach, we found that trial-by-trial reliability was moderate, with mean coefficients of variation ranging from 0.50 to 0.61 across participants and remaining consistent across stimulation intensities (Table 2). However, two-way mixed, average-measures intraclass correlation coefficients (ICC3k) demonstrated high reliability at stimulation intensities exceeding 50%, suggesting that saphenous-elicited CRs represent a viable and reliable metric for within-participant study designs, such as operant conditioning, provided stimulation intensity is sufficiently high.

### Stimulation intensity

We examined cutaneous reflex (CR) magnitudes across five stimulation intensities. Previous studies of CR EMG responses typically set stimulation intensities relative to the radiating threshold (e.g., 1.5× or 2× radiating threshold). However, for saphenous nerve stimulation, we found that these fixed ratios could exceed the pain threshold in some participants. To account for inter-individual variability, we normalized stimulation intensities for each participant based on the range between the radiating and pain thresholds. Interestingly, CRs elicited at sub-threshold intensities (−25% relative to radiating threshold) were still detectable above baseline EMG activity (Figure 5), although these responses were less reliable than those evoked at higher intensities (Table 2). Both CR magnitude and ICC3k values increased progressively with stimulation intensity, suggesting that future studies should employ relative stimulation intensities exceeding 50% to optimize reliability. Notably, CR magnitudes saturated between 75% and 100% intensity, indicating that sub-pain stimulation levels are sufficient for obtaining robust and consistent CR measurements.

### Symmetry

Finally, we assessed left–right symmetry of CR magnitudes to determine whether healthy participants exhibit lateralization of responses. All participants self-reported as right-foot dominant and the median symmetry index across participants was slightly negative (Figure 6), indicating a modest tendency toward larger right sided responses. However, at the group level, there was no conclusive evidence of systematic lateralization. For most participants, differences in left–right CR magnitudes were <10%, with a notable outlier exhibiting a difference exceeding 30%. These findings suggest that left–right CR symmetry varies across healthy individuals and is not directly related to footedness.

**Figure 6:**
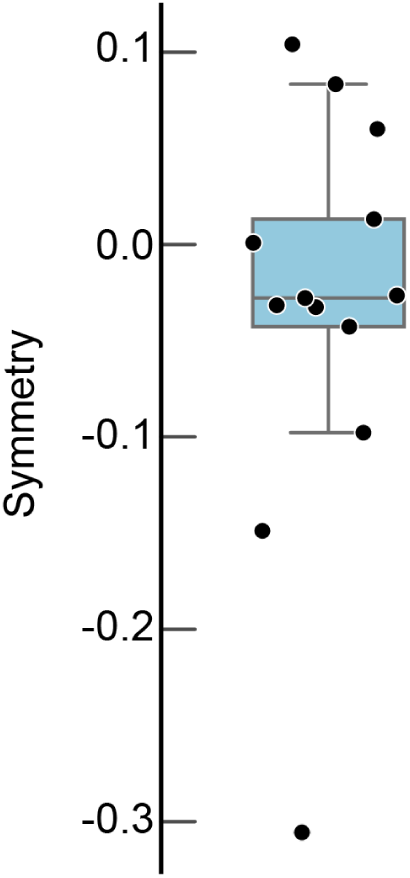
Left-right symmetry of cutaneous reflex responses. Data points denote individual participants

### Limitations

While our findings suggest that cutaneous reflexes (CRs) in the proximal leg muscles may represent a feasible target for operant conditioning, several important limitations should be considered. First, we have only assessed the test–retest reliability of CR magnitudes across multiple sessions, in a few participants to date. Although the high ICC3k values observed at higher stimulation intensities indicate that CRs can be reliably used in within-participant study designs, it remains unclear whether CR magnitudes remain stable longitudinally, across a larger population. For comparison, other evoked responses, such as the H-reflex and motor evoked potentials (MEPs), are commonly used in longitudinal studies (Manella et al., 2013; Thompson et al., 2013, 2017b, 2017a, 2018, 2019, 2022; Kim et al., 2023) yet their magnitudes can still fluctuate due to factors such as circadian rhythm (Brangaccio et al., 2024). Moreover, despite the relatively high ICC values, we observed moderate trial-to-trial variability in CR responses as indicated by the CV values (see Table 2). This variability could present challenges for operant conditioning protocols that rely on providing feedback on a per-trial basis. Whether the inherent variability in CR quantification ultimately limits the efficacy of such conditioning protocols remains an open question and warrants further investigation. Finally, when determining stimulation intensities for each participant, we relied on participant-reported sensations to determine sensory, radiating, and pain thresholds and thus cannot independently confirm that relative stimulation intensities were equal for all participants.

## Conclusion

For the first time to our knowledge, we showed that cutaneous reflexes could be effectively elicited in the proximal leg muscles via saphenous nerve stimulation. This study outlines methods and characterizations of typical side to side CR responses and their variability in healthy adults. The data here will provide a guide for typical CR responses amplitudes, latencies, thresholds, and side-to-side variability as elicited by the saphenous nerve in healthy adults. These values will serve as important guides when performing assessments and interventions on the saphenous CR responses in controlled studies with individuals in health and in disease. The knowledge and utilization of other CRs greatly expands the populations and disorders that may benefit from interventions targeting these reflexes, e.g., chronic pain disorders such as peripheral neuropathy, fibromyalgia, RSD, neuropathic pain as well as those with upright balance impairment.

